# Techno-economic assessment of animal cell-based meat

**DOI:** 10.1101/2020.09.10.292144

**Authors:** Derrick Risner, Fangzhou Li, Jason S. Fell, Sara A. Pace, Justin B. Siegel, Ilias Tagkopoulos, Edward S. Spang

**Affiliations:** Department of Food Science and Technology, University of California, Davis, CA 95616, USA; Department of Computer Science, University of California Davis, CA 95616, USA; Genome Center, University of California, Davis, CA 95616, USA; Departments of Chemistry, Biochemistry and Molecular Medicine, University of California, Davis, CA 95616, USA; Innovation Institute for Food and Health, University of California, Davis, CA 95616, USA

**Keywords:** Cell-based meat, myoblast, techno-economic assessment, bioreactor

## Abstract

Interest in animal cell-based meat (ACBM) or laboratory grown meat has been increasing, however the economic viability of these potential products has not been thoroughly vetted. Recent studies suggest monoclonal antibody production technology can be adapted for the industrialization of ACBM production. This study provides a scenario-based assessment of the projected cost per kilogram of ACBM based on cellular metabolic requirements and process/chemical engineering conventions. A sensitivity analysis of the model identified the nine most influential cost factors for ACBM production out of 67 initial parameters. The results indicate that technological performance will need to approach technical limits for ACBM to achieve profitably as a commodity. However, the model also suggests that low-volume high-value specialty products could be viable based on current technology.

**One Sentence Summary:** A model based upon cellular metabolism and engineering conventions was created to examine the economic viability of animal cell-based meat.

**Significance statement:** Animal cell-based meat (ACBM) has received a significant amount of media attention (as well as corporate investment) in recent years based on its perceived potential to displace traditional meat production, whether beef, poultry, or fish. However, a robust techno-economic assessment (TEA) of these potential products is not publicly available. Our study examined the capital and operating expenditures for potential ACBM products based upon fundamental cellular attributes, the use of proposed near-term/existing technology, and process engineering conventions. Our findings suggest that the current production pathways are far from producing cost-competitive ACBM products, as well as highlight the technical metrics that must be achieved for an ACBM product to become economically viable.

## Introduction

Global population growth and economic development are expected to double the demand for meat products by 2050 (*1*). Meanwhile, the United Nations Food and Agriculture Organization (FAO) estimates that beef and dairy cattle may be responsible for up to 5.0 gigatonnes of CO_2_-equivalent emissions, or 9% of total greenhouse gas (GHG) emissions (*2, 3*). These reported emissions are considered generalizations and a nuanced examination of an individual production system must occur to quantify CO_2_-equivalent emissions for each system (*4*). Concerns over global warming, animal welfare, and human health have prompted interest in the development of “meat alternatives” which have the organoleptic qualities of meat, but whose origin is not from a slaughtered animal (*5–8*). Analyst reports are bullish on growth in the meat alternatives sector and have predicted a significant displacement of conventional ground beef, with some reports predicting a 60-70% decrease over the next 10-20 years (*7, 8*). The predicted shift to meat alternatives would represent a disruption of a highly valuable market. In 2018, the United States processed 12.1 million tonnes of beef, including 8.5 million tonnes of retail cuts valued at US$106 billion (*9*).

Plant- and fungal-based meat alternatives are already widely available, but producers and consumers are looking to animal cell-based meat (ACBM) as the next frontier for meat alternatives. While ACBM has yet to be scaled commercially, it is currently perceived as a core component of this “2nd domestication of plants and animals”(*7*). In fact, ACBM companies have received significant early stage investments in excess of US$230 million (*10, 11*). This level of economic investment suggests the need for a rigorous assessment of the pathway to profitability for the sector.

Proposed ACBM production systems suggest existing pharmaceutical technologies could be employed for mass ACBM production (*6, 12*). Industrial-scale bioreactors would be used to proliferate myoblasts or myosatellite cells (*6, 12*). These cells then undergo a differentiation and maturation process (i.e. myogenesis) supported by a scaffolding process to form the final ACBM meat product (Fig. 1A) (*13–15*). ACBM first became a reality in 2013 with an initial public demonstration of a 140 g “hamburger” that cost over US$270,000 to produce (*16*). The high cost of production remains a significant challenge for the ACBM industry. A number of technological hurdles to lower production cost have been identified but not extensively quantified (e.g. cell senescence, high cost of growth factors, time and nutrients required for cell growth/differentiation/ maturation, and scalable scaffolding processes) (*14, 17, 18*)

**Fig. 1A.**
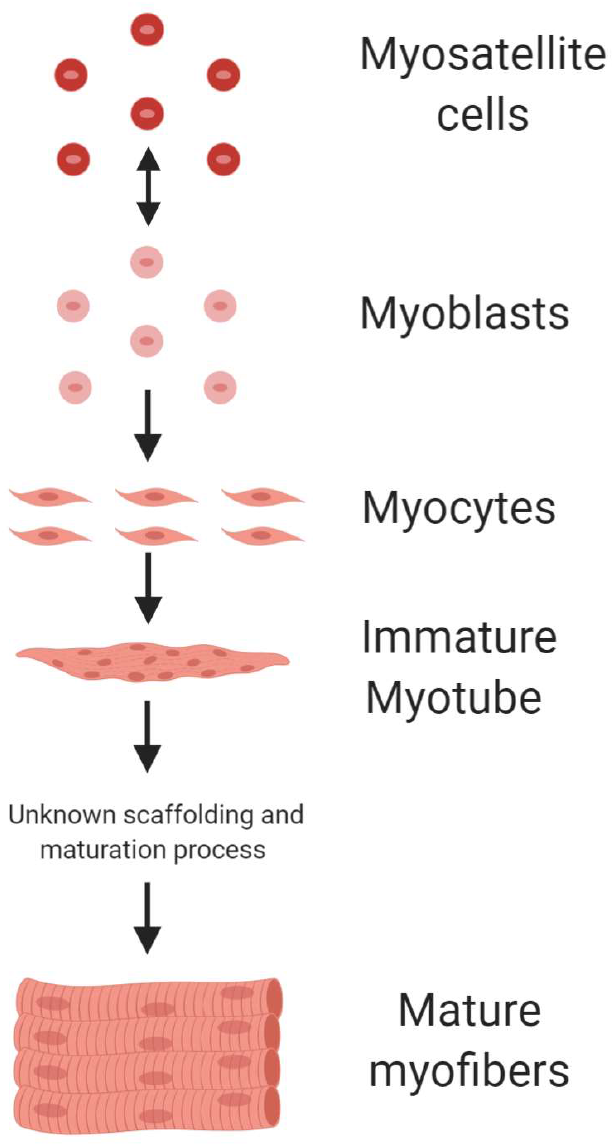
Myogenesis sequence for ACBM production.

Figure 1B illustrates a potential ACBM production system similar to monoclonal antibody production for bovine myoblasts/MSC expansion (*6, 19*). We limit our analysis here to the core bioreactor system (section “C” in Figure 1B) since industrial-scale scaffolding and maturation systems have not been defined in detail by ACBM producers. Thus, the model presented is a simplified and reduced model whose reported cost should be considered as minimum costs (Fig. 2, Eq. S1-S47, and Fig. S1-S2).

**Fig. 1B.**
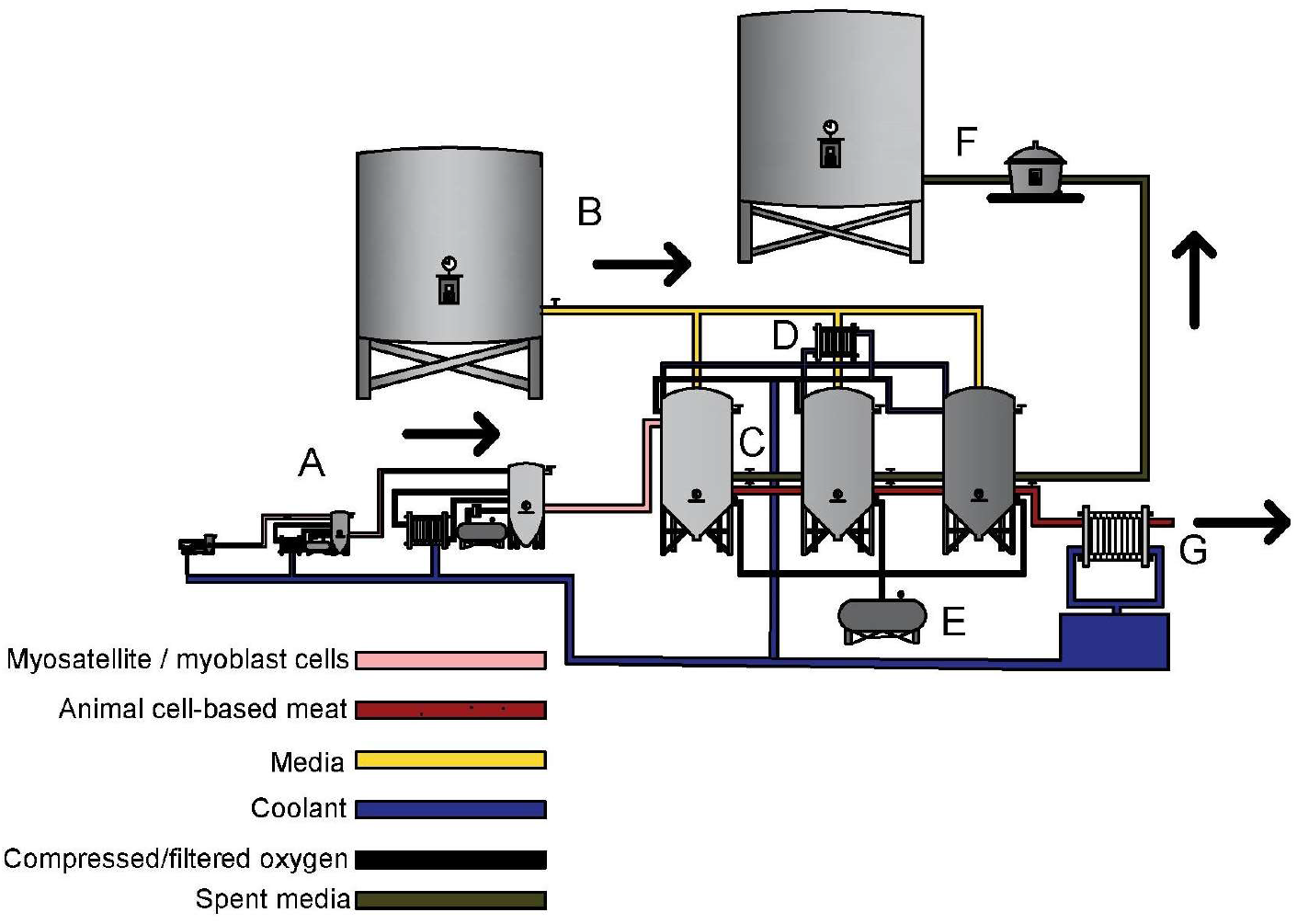
Potential industrialized ACBM production system. This system represents a potential ACBM production process without pumping system shown. A. The bioreactor seed train system 20 L, 200 L and 2000 L B. Media storage system. C. Series of 20,000 L continuous stir bioreactor system with unknown scaffolding processing occurring in bioreactor system. D. Bioreactor temperature control system. E. Oxygen supply system. F. Spent media processing system. G. ACBM cooling system. Capital expenditures only account for C, and therefore a minimum estimate of capital costs.

**Fig. 2.**
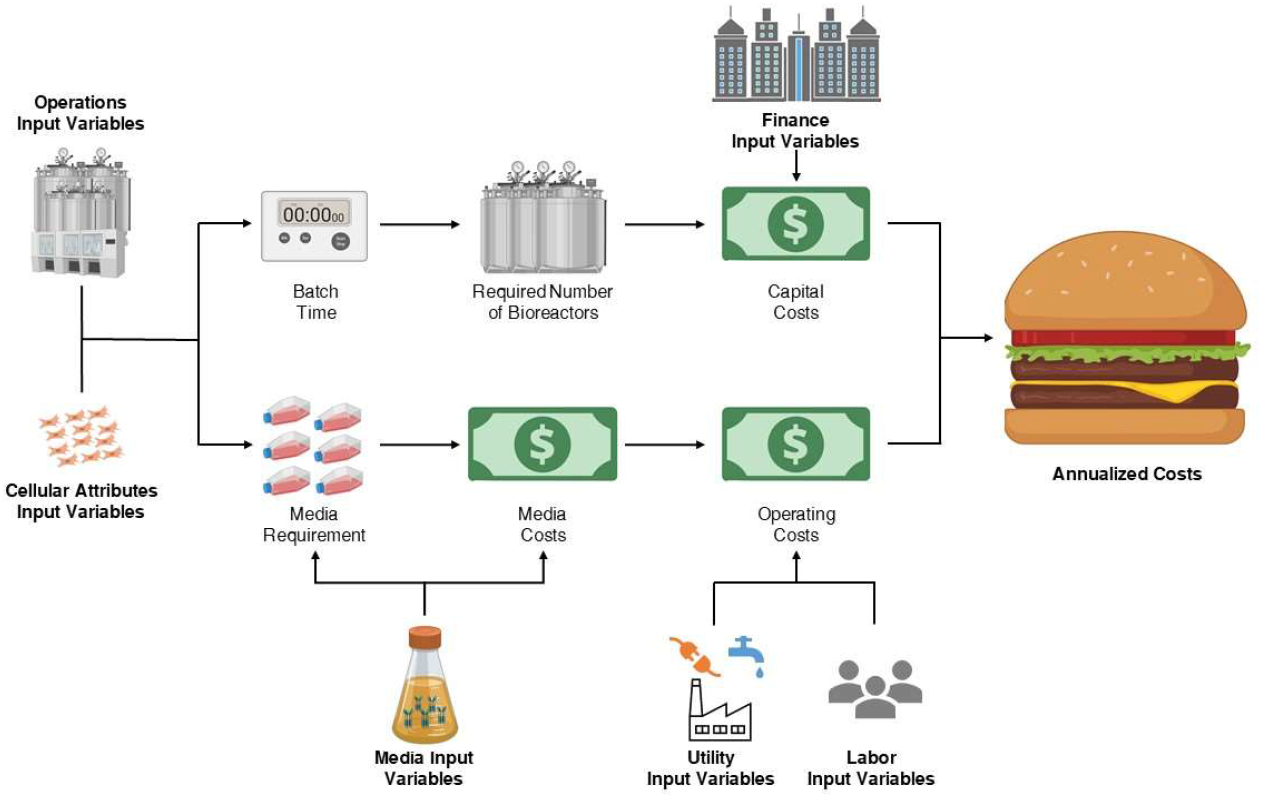
ACBM simplified economic model flow diagram. The individual input variables have been grouped into categories (Operations, Cellular attributes, Finance, Media, Utility and Labor) and can be viewed individually in Tables S1a-S1f.

Cultured bovine myoblasts using microcarriers (Cytodex® 1 or Synthemax® MC) behave similarly to human mesenchymal stem cells (HMSC) (*20*). HMSC bioprocessing is highly complex given the heterogeneity of the HMSC cultures, sensitivity of HMSC to environmental changes, spontaneous differentiation, and the necessary disassociation of cell aggregates for harvest (*21, 22*). Meanwhile, the high risk of batch contamination has led many therapeutic stem cell manufacturers to shift to single-use bioreactor systems (*22*). However, this study makes the optimistic assumption that advances in MSC/myoblast science will enable the production of MSC/myoblast using large, non-disposable, and semi-continuous bioreactor systems, and that operational issues related to bioreactor sanitation and fill rates are negligible.

## Results

Using cellular biology and chemical/process engineering conventions, we identified sixty-seven key variables that influence capital or/and annual operating costs (Tables S1a-S1f). The capital cost of a single 20 m^3^ food-grade bioreactor was estimated to be US$778,000 (*23*). We limit bioreactor size to 20 m^3^ given the sensitivity of animal cells to elevated hydrostatic pressures as compared to fungal/bacterial cells which can be viable in >500 m^3^ scale bioreactors (*24*). The annual operating expenses include fixed manufacturing costs, media, oxygen, energy, process water, and wastewater treatment costs.

To understand the impact of each model variable on the estimated capital and annual operating expenses, we performed a robust sensitivity analysis (fig. 3 and Table S2). We applied six global sensitivity analysis algorithms (Derivative-based Global Sensitivity Measure, Delta Moment-Independent Measure, Morris Method, Sobol Sensitivity Analysis, Fourier Amplitude Sensitivity Analysis, Random Balance Designs-Fourier Amplitude Sensitivity Test) to identify the top nine factors that most influenced capital and annual operating expenses by consolidating the top 5 parameters across all six algorithms. These nine factors were then clustered into technological components (including maturation time, fibroblast growth factor 2 (FGF-2) concentration and costs, glucose concentration, glucose consumption rates, oxygen consumption rate and transforming growth factor beta (TGF-β)) and cell-based components (e.g. average cell volume and density).

**Fig. 3.**
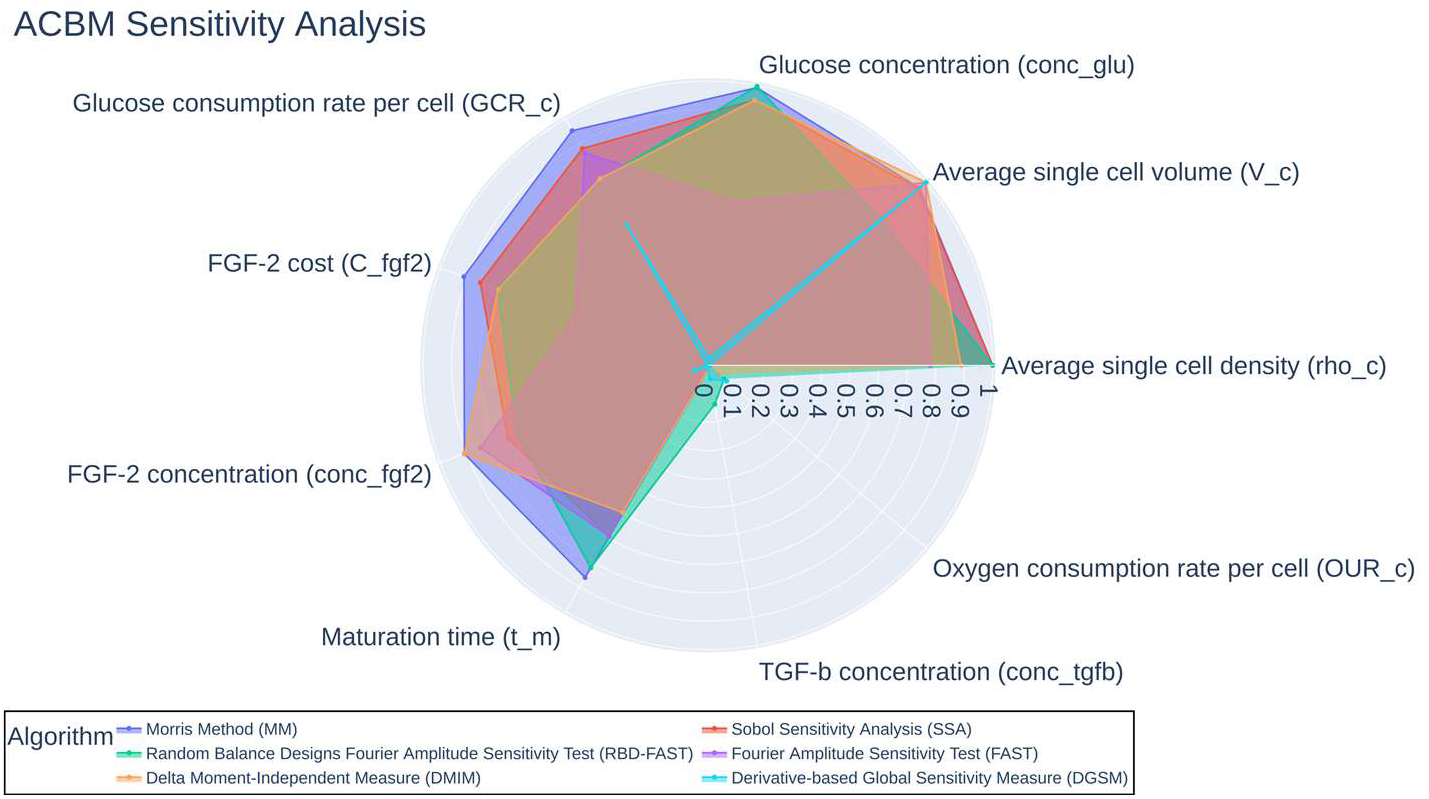
ACBM sensitivity analysis of key model variables. Each algorithm independently examined 67 parameters for sensitivity. The 5 parameters exhibiting the most sensitivity were selected from each algorithm. This resulted in 9 unique parameters visualized in the figure. The sensitivity measurements of the algorithms were scaled from 0 to 1 using minimum-maximum normalization except DGSM. The measurement of DGSM was first scaled by taking its sixteenth root and then normalized from 0 to 1 by minimum-maximum. Abbreviations for each of the 9 unique parameters are provided for reference to the input variables in Data S1

The results from the sensitivity analysis then informed the specification of four technology development scenarios (Tables 1 and S1a-S1f). Scenario 1 represents a baseline scenario a based on existing ACBM production, including 2019 cost estimates for animal serum-free media and growth factors (*12*). Scenario 4 was designed as a bookend scenario, where nearly all technical challenges are resolved, including reduced growth factor costs, increased MSC/myoblast tolerance to glucose concentrations, decreased MSC/myoblast doubling and maturation time, and reduced basal media costs (*6, 12, 14, 17*). Scenario 2 represents a mid-point scenario between Scenarios 1 and 4, and Scenario 3 adapts Scenario 2 by eliminating FGF-2 growth factor costs. To incorporate economic scalability, we also examined the capital and annual operating expenditures to produce enough ACBM to replace 1% of the United States beef market (121,000,000 kg) (*9*).

The results of our calculations indicate that ACBM production will only approach economic viability as a commodity when the significant technical challenges are overcome as outlined in Scenario 4 (Table 1). In Scenario 1, the cost per kilogram remains exceedingly expensive at approximately US$400,000. Scenarios 2 and 3 illustrates the significant impact of reducing the cost of FGF-2, which reduces the operating cost of ACBM by an order of magnitude from Scenario 1. Only in Scenario 4 does ACBM approach commodity level prices at approximately US$2 per kg.

**Table 1.**
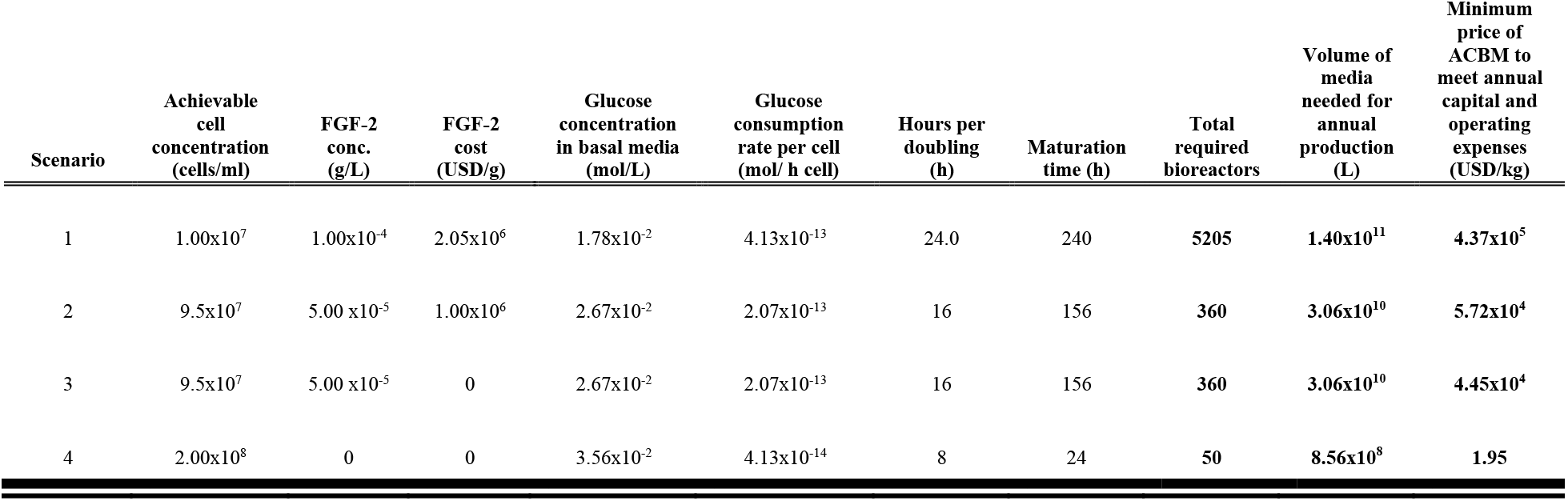
Model scenarios and results for replacing the equivalent of one percent (121,000,000 kg/year) of United States beef production with ACBM. Columns 2-8 are scenario parameters and bolded numbers represent model results. These should be considered minimum production inputs and expenditures. The model only examines main bioreactor equipment costs and some but not all operating costs. Model limitations can be found in the supplemental material. Current commodity beef prices are approximately US$2 per kg.

| Scenario | Achievable cell concentration (cells/ml) | FGF-2 conc. (g/L) | FGF-2 cost (USD/g) | Glucose concentration in basal media (mol/L) | Glucose consumption rate per cell (mol/ h cell) | Hours per doubling (h) | Maturation time (h) | Total required bioreactors | Volume of media needed for annual production (L) | Minimum price of ACBM to meet annual capital and operating expenses (USD/kg) |
| --- | --- | --- | --- | --- | --- | --- | --- | --- | --- | --- |
| 1 | 1.00x10<sup>7</sup> | 1.00x10<sup>-4</sup> | 2.05x10<sup>6</sup> | 1.78x10<sup>-2</sup> | 4.13x10<sup>-13</sup> | 24.0 | 240 | **5205** | **1.40x10<sup>11</sup>** | **4.37x10<sup>5</sup>** |
| 2 | 9.5x10<sup>7</sup> | 5.00 x10<sup>-5</sup> | 1.00x10<sup>6</sup> | 2.67x10<sup>-2</sup> | 2.07x10<sup>-13</sup> | 16 | 156 | **360** | **3.06x10<sup>10</sup>** | **5.72x10<sup>4</sup>** |
| 3 | 9.5x10<sup>7</sup> | 5.00 x10<sup>-5</sup> | 0 | 2.67x10<sup>-2</sup> | 2.07x10<sup>-13</sup> | 16 | 156 | **360** | **3.06x10<sup>10</sup>** | **4.45x10<sup>4</sup>** |
| 4 | 2.00x10<sup>8</sup> | 0 | 0 | 3.56x10<sup>-2</sup> | 4.13x10<sup>-14</sup> | 8 | 24 | **50** | **8.56x10<sup>8</sup>** | **1.95** |

The cost of the bioreactor was the main driver of capital costs in the model. To displace the demand for beef in the U.S. by 1%, the scenarios ranged from requiring the deployment of 5205 to 50 bioreactors (20 m^3^) at a total capital cost of 4 billion to 37 million U.S. dollars. The capital expenditures in scenario 3 remain the same as scenario 2 since eliminating the growth factor cost has no impact on the capital expenditures. Finally, it is important to reiterate that these costs are based on estimates for standard food-grade bioreactors and that more sophisticated bioreactors (i.e. single-use or novel perfusion systems) may substantially increase capital costs.

While capital expenditures are significant, the operating expenses (largely based upon cellular metabolism and media consumption) represent a substantial hurdle for the large-scale production of ACBM. Achieving the outcomes presented in Scenario 4 would require significant technological advancements on multiple fronts as specified by the model, where media costs are reduced from 376.80 US$/L to 0.24 US$/L, glucose/media consumption is reduced by an order of magnitude, and cell growth and maturation times are heavily decreased from 24h to 8h and 240h to 24h, respectively.

## Discussion

The results of the scenario analysis clearly highlight and quantify the technological and economic challenges for ACBM to reach commercial viability. We suggest the following three areas of focus to reach techno-economic feasibility, which we will discuss further: cell selection or engineering to lower the media consumption rate, reducing or eliminating the cost of growth factors, and scaling up of perfusion bioreactors.

The analysis identified cell metabolism as a key limiting factor for the economic viability of ACBM, so understanding and potentially manipulating cellular metabolism represents a key area of innovation for driving down operating costs. The glucose consumption rate of cultured cells establishes the media requirements in our model, which is by far the largest operating expense for ACBM production. Scenario 1 was based upon reported human embryonic stem cells’ glucose uptake rate. These cells were likely exhibiting a Warburg metabolism (aerobic glycolysis) based upon their lactate production rates (*25, 26*). This metabolic mode is common during cell proliferation; however, it is energetically less efficient than oxidative phosphorylation (i.e. production of 2 ATP vs. a theoretical 38 ATP per glucose molecule) (*26*). Engineering and/or screening for cell lines which shift rapidly from a Warburg metabolism to a more glucose-efficient metabolism represents an opportunity to reduce the media consumption rate in line with Scenario 4.

In healthy cells, glucose uptake is stimulated by growth factors such as insulin, FGF, or/and TGF(*25, 27*). Our model highlights that growth factors are a major contributor to ACBM production expenses, with FGF-2 being particularly impactful. Thus, eliminating the need for FGF-2 would significantly reduce costs. One potential pathway for this solution would be to leverage the ability of cancer cells to increase glucose uptake rates and exhibit cell proliferation without the presence of growth factors (*25*). Thus, cell lines could be engineered or identified to express oncogenes related to these traits. However, utilizing cultured cells that behave similar to cancer cells would likely be very challenging from both a regulatory perspective as well as for consumer acceptance.

Our model indicates that cellular metabolic requirements will require multiple changes of media per batch and higher cell concentrations (*28, 29*). The use of perfusion bioreactors could deliver these capabilities for ACBM production (*30*). Concentrations of 2.0 x10^8^ cells/ml have been reported for Chinese hamster ovary (CHO) cells in a lab-scale, disposable perfusion bioreactor system (*30*). However, this is a profoundly different technology than the large-scale, continuously stirred bioreactors we assume in our model. To the authors’ knowledge, a perfusion bioreactor system with a 20 m^3^ working capacity is not currently in existence for myoblasts/MSC propagation.

ACBM has been presented as a potentially disruptive technology that can transform the global meat sector. However, our techno-economic analysis of this alternative meat production pathway suggests that the profitable mass production of products composed entirely of ACBM remains a significant challenge. Our model indicates that several technical challenges must be overcome before industrial scale-up is likely to be profitable. Media consumption rates must be measured and optimized at the cellular level and the costs of growth factors must be significantly reduced or eliminated altogether.

While these factors indicate that ACBM may not be economically viable as a commodity for some time, it does not preclude the potential to enter the market place sooner as a minor ingredient which lends desirable organoleptic qualities to an otherwise plant-based product. Alternatively, there may be opportunity for viable competition in the specialty foods markets, where ACBM costs compare more favorably to such items as almas beluga caviar (US$10,000/kg), Atlantic bluefin tuna (US$6,500/kg), and foie gras (US$1,232/kg) (*31*).

Our model has highlighted some of the significant economic challenges which impede the techno-economic viability of ACBM, but it is not comprehensive. Given the uncertainty of ACBM production, our model should be considered a starting point for those interested in the scalability of ACBM. To enable further, and customizable, exploration of how advances in technology might inform ACBM production costs, we have developed an open-source, webbased version of our model that is publicly available at http://iifh-meat-cost-calculator.s3-website-us-west-2.amazonaws.com/].

## Materials and Methods

To determine the economic viability of animal cell-based meat (ACBM), we developed a model using standard process and chemical engineering methods. The model system is a semi-continuous-batch production system operating at capacity year around and does not account for fill times, sanitation between batches or any operational downtime. Table S3 provides a list of equipment that would likely be necessary for industrial ACBM production. The costs were broadly broken down into annual operating costs and capital expenditures then annualized. All equations and variables are available in the equation and variable lists in the supplementary material as well as in the python code associated with our model.

### Capital Expenditures of an ACBM plant

We accounted for the volume each myoblast/myosatellite cell (MSC) occupies with the operating constraint that the total cell volume cannot exceed bioreactor operating capacity for each batch. Cell volumes are variable, so a reported volume estimate of 5 × 10^−15^ m^3^ cell^−1^ was used (*12*). Eukaryotic muscle cell density is approximately 1060 kg m^−3^ and was used to estimate mass of ACBM per batch (*32*). The actual density of ACBM may be lower due to incorporation of bovine adipose cells or other sources of fat. A decrease or increase in batch time influences economic viability of ACBM production. The batch time is the sum of the cell growth phase and maturation time (equation 1). The cell concentration is considered a variable that can change with technological innovation. Using a given cell concentration, the mass of each batch of ACBM was determined using equations 2-4. The batch time was then used to calculate the annual ACBM batches per bioreactor and the number of bioreactors required to achieve the desired annual ACBM production mass (equations 5 and 6).

Cost estimates of food-grade bioreactors were calculate using a method which accounts for equipment scaling, installation, and inflation (equations 7 and 8) (*23*). This method applies a set unit cost of $50,000 m^−3^ for a food grade bioreactor and a common scaling factor of 0.6 (*23*). To account for inflation and changes in cost over time the Chemical Engineering Plant Cost Index (CEPCI) values for heat exchangers and tanks were used to determine an adjusted value factor (*33, 34*). Adjusted value factor of 1.29 was determined dividing the recent CEPCI values with the values from when the set unit cost was referenced. The Lang factor is used to estimate cost associated with installation and piping. This factor can range from 1.35-2.75 for traditional food production operations and to up to 4.80 for fluids processing (*35*). A Lang factor was estimated to be 2 for all scenarios. For new plant cost the Lang factor value should be increased by 1 (*35*). This estimated the minimum capital expenditures for the required number of bioreactors which are necessary to meet the desired ACBM production mass. This method doesn’t account for any other equipment which would likely be necessary for ACBM production (Table S3) besides the primary bioreactor systems.

### Operating costs of an ACBM plant

The potential manufacturing cost of an ACBM plant can be broken into three categories: Fixed manufacturing costs, variable capital costs and indirect (overhead) costs. All fixed manufacturing costs were estimated as a percentage of the fixed equipment costs except loan and equity interest (equation 9) (*35*). These costs include equipment maintenance, insurance, taxes and royalties costs (*35*). Indirect costs are not accounted for in our model since these costs are outside of plant operation expenses and will vary company to company. Our model provides an estimate of several variable capital costs related to downstream ACBM production. Costs associated with general meat production such as packaging material and facility lighting are not included. The variable costs estimated in our model include ingredients, raw materials, utilities and labor costs. Equation 10 accounts for all the operating costs associated with the model we have provided.

### Ingredients and raw materials

A key material for animal serum-free ACBM production is the specialized media required for myoblasts/MSCs growth. Our model examines the use of Essential 8, an animal free growth medium which contains over 50 ingredients including ascorbic acid 2-phosphate, sodium bicarbonate, sodium selenite, insulin, transferrin, fibroblast growth factor-2 (FGF-2), and transforming growth factor beta (TGF-b§) (*12*). A report from the Good Food Institute provides an excellent breakdown of the individual components of Essential 8 media and the 2019 pricing of each media component (*12*). Cell glucose consumption rates can vary based upon several factors including glucose concentration present in the growth medium and the metabolic pathways being utilized by the cell (*36, 37*). Glucose consumption rates have been reported to be between 2 to 20 nmol^1^ million cell^−1^ min^−1^ in human stem cells (*37*). While there can be many limiting factors in a complex medium system; glucose consumption and the total number of cells in the bioreactor were used to estimate the media requirements and expense per batch. The starting glucose concentration is reported to be 1.78×10^−2^ mol L^−1^ (*12*). Only media used in the main bioreactor was accounted for. An oxygen supply is also critical for aerobic cell culture and is also considered an operating expense for ACBM production. Equation 11 was used to determine total amount of myoblasts/MSCs in the bioreactor at a given time. During the growth phase, the glucose consumption rate changes as time changes and this was accounted for using equation 12. The total glucose required for the growth phase was determined by equation 13. The total glucose required per batch was determined by adding the total glucose used in the maturation and growth phase (equations 14 and 15).

The media requirement was then determined by examining the total amount of glucose in the Essential 8 media. To understand the volume requirement per batch, a charge was deemed the equivalent to the working volume of the bioreactor. This assumption was done to account for any innovations related to vascularization and does not account for the volume of the cells. The total media volume required per batch/year and total annual media costs were determined by equations 16-19.

An oxygen supply is critical for aerobic cell cultures and is also considered an operating expense for ACBM production. The oxygen levels in the bioreactor were assumed to be kept in a steady state concentration of 2% for optimal cell growth (*21, 38*). This is expressed by equation 20 (*39*). The initial oxygen needed for the bioreactor system was determined by equation 21. The annual oxygen requirement was determined in the same manner as the media requirement and is calculated using equations 11 and 22-27.

### Utility related expenses

Our model accounts for some bioreactor operating expenses. These should be viewed as theoretical minimum estimates based upon conventional thermodynamic equations. The energy requirements for heating the media, cooling the bioreactors and cooling of the ACBM mass leaving the bioreactor systems were estimated. The water/media was assumed to enter the facility at approximately 20 °C. The media is also assumed to have an isochoric specific heat of approximately water. The density of the media was assumed to be 1 kg L^−1^ and would be heated to 37 °C. The minimum energy required to heat the media was calculated using equation 28. The metabolic consumption of glucose and oxygen produces heat which must be removed from the system. Approximately 470 kJ of heat is released per mol of O_2_ consumed during glucose combustion (equation 29) and this value was used to approximate cellular heat generation (*39*). The minimum energy required to be removed from the system to ensure cell health was calculated using equation 30. The ACBM mass leaving the bioreactor must be cooled from 37 °C to 4 °C to ensure food safety standards are maintained (*40*). The specific heat of ACBM is assumed to be the same as beef which is 2.24 kJ kg^−1^ °C^−1^ (*41*). An estimation of energy used during the cooling process (equation 31) was made based on the efficiency of the heat exchanger system.

Energy costs can be variable depending upon the location, time of day and amount used. A yearly national grid average for industrial electricity and natural gas prices was obtained from the United States Energy Information Administration (EIA) from 1999-2019 (*42*). One thousand cubic feet of natural gas contains approximately 303.6 kWh of potential energy and the cost per kWh was determined using this value (*43*). The average costs were normalized to January 2019 prices using the CPI inflation calculator (Table S4 and S5) (*44*). To estimate the energy/electricity cost a comparison of the industrial price of natural gas and electricity was made from 1999-2019 (Fig. S1). Equation 32 was derived from a linear relationship of the cost of electricity and natural gas (Fig. S2). Equation 32 was then used to estimate energy/electricity costs from a public supplier. Natural gas was chosen since it is the most used source of energy for electricity production in the United States in 2019 (*45*). The costs of energy/electricity produced via an onsite boiler-turbine system was estimated by equation 33. A steam pressure of 42.5 bar is assumed because it is used as a reference pressure for cost of steam production and is adequate for steam turbine electricity production (*46, 47*). Solar generation of electricity was considered as well and was estimated to have a negligible operating cost for the facility. The equipment costs for solar are not accounted for since this is a facility dependent item. Equation 34 estimates the minimum cost of energy at an ACBM production facility.

Our model assumes media will be produced onsite given the scale of the operation. All water used for media production is considered process water, however it should be noted that deionized water could be required due to the operational sensitivity of myoblasts/MSCs. Compressed air is a common utility in food production facilities; however, it is not estimated in this analysis due to being a site-specific consideration. Cost of sterile filtration of the water for media production is not accounted for. The spent media is considered wastewater and must be treated to comply with environmental regulations (*48*). The wastewater is assumed to be treated by a filtration and biological oxidation step. Cost estimates have been made for process water and wastewater treatment and these estimates have been adjusted to January 2019 values to account for inflation (Table S6) (*44, 46*). It should be noted that this does not account for water used for sanitation or for losses during the production process. Equation 35 is used to estimate the annual process and wastewater costs.

### Labor related expenses

Our scenarios assume that the ACBM production facility is operating 24 hours/day and year around. It is assumed the facility is fully staffed and no overtime is required. Each shift is assumed to be an 8-hour shift. The facility is also assumed to be in the United States in an area of standard income. The required production operators (required manpower) for the ACBM production facility per shift is estimated by amount and type of processing equipment in the facility (Table S3) (*23, 46*). This processing equipment could include centrifugal pumps, plate filters, media holding vessels, heat exchangers, bioreactor seed train, positive displacement pumps and bioreactors. In the four scenarios, this equipment was deemed site specific and only the main bioreactors were accounted for. The labor cost were determined using the mean hourly rate, $13.68 (USD h^−1^) for a meat packer (*49*). The labor costs were estimated using a factorial method with a labor cost correction factor (equations 36-38) (*46*).

### Finance related expenses

Our model accounts for the expenses related to equity recovery and debt using a standard finance calculation (equations 39-46) (*50*). For all scenarios, the input variables were kept constant. Equations 39-46 convert the capital expenditures to an annual cost which is used to calculate the total annual minimum costs in conjunction with the annual operating costs (equation 47).

### Sensitivity analysis

We performed a sensitivity analysis of the ACBM price model using 6 algorithms that use different approach to variance and rate of change to assess sensitivity: the Derivative-based Global Sensitivity Measure (DGSM), Delta Moment-Independent Measure (DMIM), Morris Method (MM), Sobol Sensitivity Analysis (SSA), Fourier Amplitude Sensitivity Test (FAST), and the Random Balance Designs Fourier Amplitude Sensitivity Test (RBD-FAST). We used the SALib Python package for this work (*51*). Additional information regarding sensitivity analysis algorithms can be found in the supplementary material.

## Author contributions

Derrick Risner contributed to the conceptualization, methodology, investigation, writing (original and final draft preparation, review and editing) and visualization. Fangzhou Li contributed to the formal analysis, writing (original and review) and visualization. Jason Fell contributed to the software development, visualization and writing (review). Sara Pace contributed to methodology, visualization, writing (review), and validation. Justin Siegel contributed to the conceptualization, methodology, writing (review and editing), supervision, project administration and funding acquisition Ilias Tagkopoulos contributed to the supervision, formal analysis, writing (review) and visualization. Edward Spang contributed to the conceptualization, methodology, writing (review and editing), supervision, project administration and funding acquisition.

