## Supplementary material for "Techno-economic assessment of animal cell-based meat"

**Other Supplementary Materials for this manuscript includes the following:**

Code available upon reviewer request and will be made public upon publication.

Data S1. Techno-economic analysis and sensitivity analysis python code for ACBM
https://github.com/IBPA/IBPA-Collection-of-Reproducible-Code-and-Results/tree/master/2020_Artificial_Meat
Data S2. Techno-economic analysis web-based program for ACBM
http://iifh-meat-cost-calculator.s3-website-us-west-2.amazonaws.com/

**Supplementary Text**

<u>Model limitations</u>

In human pluripotent stem cells, as the cells exit pluripotency and enter the initial differentiation phase a metabolic shift to mitochondrial OXP occurs (*1*, *2*). A similar shift occurs as myoblasts fuse differentiate into myotubes (*3*). As myoblasts differentiate into myotubes it has been reported that the metabolic rate is maintained despite a greater reliance on OXP pathway for ATP production(*3*, *4*). However, it is not known if this metabolic rate will be maintained during the undefined scaffolding and maturation process. During this undefined scaffolding and maturation process, the myotubes diameter could potentially increase 20-fold(*5*–*7*). Our model assumes glucose and oxygen uptake rate are maintained during this process; however, these values could change to meet the metabolic needs of the maturing myotubes. Once the myotubes mature, they rely upon OXP to meet their metabolic needs and this shift may require an adjustment to operation factors such as an increased or decreased media or oxygen supply.

Our model did not account for amino acid uptake rates due to glucose being the most consumed nutrient in cell culture, however amino acid (AA) metabolism should be a consideration for commercial scale up. An example of the importance of this consideration is that stem cell amino acid metabolism can vary species to species (*8*, *9*). Bovine and mouse embryonic stem cells are sensitive to extrinsic deprivation of threonine, whereas human embryonic stem cells are not sensitive extrinsic deprivation of threonine, but require increased levels of methionine (*9*–*11*). This extrinsic threonine requirement does not apply to other mouse or bovine cells which are proliferating(*8*). This illustrates how these requirements can vary by species and by cell type.

Glutamine is utilized as both a nitrogen donor and energy substrate in proliferating myosatellite/myoblast cells (*12*, *13*). Glutamine is the second most consumed nutrient in animal cell cultures and contributes to nucleic acid, protein and lipid production (*14*). Glutamine concentration has been show to influence the myoblasts proliferation rate with 300 μM being reported as the optimal conditions for human myoblasts proliferation (*13*). This indicates that amino acid levels in the media could potentially influence operating costs via increased or decreased doubling times. This would likely be cell line dependent and should again be a consideration for companies wishing to develop multiple products from different cell lines.

The volume of animal cells also plays an important factor in our modeling which accounts for the volume of each cell. Animal myoblasts cells volume are orders of magnitude larger than common prokaryotic or single cell fungi (*15*). This places hard constraints on the number of cells a single bioreactor can produce per batch i.e. bioreactor with a working volume of 20 $m^3$ can only produce the number of cells whose total volume is $20m^3$. This does not account for repulsive forces or for the media within bioreactor. While this was done to account for any innovations in vascularization it makes the model less conservative and should be a consideration for any company considering scale up. It also does not account for cellular volume increases during the unknown scaffolding and maturation phase. The diameter of the myotube can increase up to 20 times it's original size as contractile protein is formed (*5*–*7*). This increase in size of the cells during maturation could make the bioreactor more efficient, however it was not included in our model due to the unspecified nature of the commercial process.

Figure 2B represents a potential upstream production system for ACBM, however the capital expenditures that were estimated by our model only estimate the cost of a series of 20,000 L continuous stirred bioreactors designated by letter A. We did not adjust the maximum bioreactor operating capacity of the bioreactors in any scenario due to fragility of animal cells which lack a cell wall and cannot withstand the hydrostatic pressures which yeast or prokaryotic organisms can (*16*). Innovations in bioreactor design could potentially increase the maximum working capacity. An increase in bioreactor working capacity would potentially lower capital expenses and annual operating costs. However, this would initially increase the base cost ($50,000/m$^3$) of the bioreactor measured in our model. In a more detailed analysis as the metrics we have outlined are achieved, interest rate and learning curve equations could be applied to estimate capital and operating expenses in finer granularity. We also assume that the unknown scaffolding and maturation process could be accomplished within the bioreactors. If a separate bioreactor or maturation vessel is needed this would also increase capital expenditures. We did not account for the other equipment since this will be a site-specific variable. The Lang factor is used to estimate actual cost of equipment by accounting for installation related expense. A Lang factor of 2 was chosen for all scenarios to represent a food/bioprocessing facility that could be easily configured to accommodate ACBM production. However, a Lang factor of 2 is considered to be low by general conventions for a brand new facility or novel technology; a Lang factor of 3 to 5 would be more appropriate (*17*). We anticipated that once the ACBM is cooled it will be processed in a manner similar to other ground meat products. We also did not account for any additional ingredients being added to the product. Cellular propagation technology could potentially be applied for myoblasts/MSC propagation. Cytodex® 1 microcarriers have been employed for bovine myoblasts proliferation and achieved a cell concentration of approximately $9 \times 10^6$ cells/ml (*18*). Our model does not account for this technology or any additional propagation technology which may increase capital or operating expenses. It has also been reported that bovine muscle satellite cells have been cultured with hemoglobin and myoglobin(*19*). Costs associated with additional ingredients or media supplementation have not been accounted for and could substantially increase the annual operating expenses.

Additional sensitivity analysis information

All sensitivity analysis calculations were conducted using the SALib Python package (*20*). Regarding sampling techniques and parameters, Delta Moment-Independent Measure (*21*, *22*) and Random Balance Designs Fourier Amplitude Sensitivity Test (*23–25*) used 1000 samples generated using Latin hypercube sampling (*26*), where Random Balance Designs Fourier Amplitude Sensitivity Test used the inference number of 10. Sobol Sensitivity Analysis used 1000 samples generated using Saltelli sampling (*27–29*). Morris Method was sampled with 1000 trajectories and 4 grid levels (*30*). Fourier Amplitude Sensitivity Test used 1000 samples with the inference number of 4 (*31*). Derivative-based Global Sensitivity Measure used 1000 samples with finite difference step size of 0.0001 (*32*). The result of the sensitivity analysis is shown in Figure 3 and table S2.

Variables list

Variables are listed in the order they appear in the equations.

$t_b$= time of batch (h)
$t_{gf}$= Time growth phase ends (h)
$t_m$= Time of maturation phase (h)
$F_c$= Final concentration of cells in bioreactor (cells $L^{-1}$)
$B_v$= Bioreactor working volume (L)
$N_c$= Total number of cells in bioreactor (cells)
$V_c$= Volume of single cell ($m^3$ $cell^{-1}$)
$V$= Volume ($m^3$)
$\rho_c$= Density of muscle cell (kg $m^3$)
$M_b$= mass of ACBM produced per batch (kg $batch^{-1}$)
$b_{BY}$= Number of batches a single bioreactor can produce in year (batches $year^{-1}$)
$M_{BY}$= Mass of ACBM a bioreactor can produce in a year (kg $year^{-1}$)
$M_{DY}$= Desired annual mass of ABCM (kg)
$B_T$ = Total number of bioreactors required to annual production goal
$C_{eq}$= Total equipment costs (USD)
$C_F$= Fixed equipment cost (USD)
$f_{Aj}$= Adjusted value factor for equipment j
$C_{Uj}$ = Unit costs for equipment j
$U_j$= Base unit for equipment j
$U_{aj}$= Actual unit for equipment j
$f_s$= Scale factor for equipment j
$f_L$= Lang factor
$f_{FM}$= Fixed manufacturing cost factor
$C_{FM}$= Fixed manufacturing costs (USD)
$C_{op}$= Annual operating costs (USD)
$C_{mY}$= Total annual costs of media (USD)
$C_{O_2Y}$= Total annual costs of oxygen (USD)
$E_{Hm}$ = Minimum energy required to heat media (kWh)
$E_{BR}$ = Minimum energy required bioreactor heat removal (kWh)
$E_{ACBMR}$= Minimum annual energy required for ACBM heat removal (kWh)
$C_L$= Estimated annual labor costs (USD)
$C_E$= Cost of energy (cents $kWh^{-1}$)
$C_W$= Annual process water and wastewater costs (USD)
$c_t$ = Total number of cells at time (t)
$c_o$ = Total number of cells present in inoculum (cells)
$t_D$ Doubling time (h)
$t$ = Time (h)
$GCR_B$ = Glucose consumption rate within the bioreactor (mol $h^{-1}$)
$GCR_c$= Glucose consumption rate per cell (mol $h^{-1}$ $cell^{-1}$)
$G_{Gg}$=Total moles of glucose required for growth phase (mol)
$G_{GM}$= Total moles of glucose required for maturation phase (mol)

$G_G$= Total moles of glucose required per batch (mol)
$m_{ch}$= Total media charges per batch (charge)
$M_{Gch}$= Moles of glucose per charge (g)
$V_b$= Total volume of media required per batch (L)
$V_{ch}$= Volume of charge or bioreactor (L)
$V_m$ = Total media volume per year (L year$^{-1}$)
$b_y$ = Batches per year
$C_{mL}$=Cost of media per liter (USD L$^{-1}$)
$OUR_B$ = Oxygen uptake rate in bioreactor (mol s$^{-1}$)
$OTR_B$ = Oxygen transfer rate in bioreactor (mol s$^{-1}$)
$k$ = mass transfer coefficient (m s$^{-1}$)
$A$ = mean bubble specific interfacial surface area (m$^2$)
$e_{con}$= equilibrium concentration (mol m$^{-3}$)
$a_{con}$= actual dissolved oxygen concentration (mol m$^{-3}$)
$O_2^i$ = Initial oxygen in required in the system (mol)
$\rho_m$ = Density of media (kg L$^{-1}$)
$P_{O_2}$ = Percentage of oxygen ($O_2$) in media by weight (%)
$O_2^{mol}$ = molar mass of $O_2$ (kg mol$^{-1}$)
$OUR_c$= rate of oxygen consumption per cell mol cell$^{-1}$ h$^{-1}$
$O_2^g$ = Total oxygen required for growth phase per batch (mol)
$O_2^M$=Total oxygen required for maturation phase per batch (mol)
$O_2^b$= Total oxygen used per ACBM batch (mol)
$O_2$ = Total amount of oxygen required per year (mol)
$C_{O_{2Y}}$ = Total annual costs of oxygen (USD)
$C_{O_2}$ = Cost of oxygen (USD mol$^{-1}$)
$M_{mY}$=Mass of media used per year (kg)
$\Delta T$ = Temperature difference (˚C)
$W_{C_v}$= Specific heat of water at constant volume (kWh kg$^{-1}$ ˚C$^{-1}$)
$\in_{Hm}$ = Energy efficiency of heating system (%)
$O_2$ = Oxygen required annually (mol)
$h$ = Heat released per mol of oxygen consumed (kWh mol$^{-1}$)
$\in_{BR}$ = Energy efficiency of bioreactor cooling system (%)
$ACBM_{C_v}$ = Specific heat of ACBM (kWh kg$^{-1}$ ˚C$^{-1}$)
$\in_{ACBMR}$ = Energy efficiency of ACBM cooling system (%)
$C_{EP}$ = Cost of electricity from a public supplier (USD kWh$^{-1}$)
$C_{NG}$= Cost of natural gas (USD 1000 ft$^{-3}$)
$C_{bT}$ = Cost of energy from onsite boiler-turbine system (USD kWh$^{-1}$)
$C_{NGP}$ = natural gas price (USD kWh$^{-1}$)
$\epsilon_{bT}$ = boiler-turbine system efficiency (%)
$f_{EP}$ = percentage of electricity produced by from a public supplier (%)
$f_{bT}$ = percentage of energy produced by on site boiler-turbine system (%)
$C_{PW}$ = Process water costs (USD m$^{-3}$)
$C_{WF}$ = Wastewater filtration costs (USD m$^{-3}$)
$C_{BO}$= Biological oxidation of wastewater costs (USD m$^{-3}$)
$P$ = required manpower (production workers)

$P_j$ = production worker required for single piece of equipment
$j$ = Individual piece of equipment
$N$ = All downstream equipment used in downstream ACBM production
$f_{lab}$ = Labor cost correction factor
$f_C$ = Country effect
$f_{Sca}$ = Supervising and clerical assistance
$f_T$ = Advanced technological and automating
$f_Q$ = Skilled and qualified level of the personnel
$f_B$ = Social benefits
$f_O$ = Overtime work
$C_{Lab}$ = Estimated annual labor costs (USD)
$t_y$ = Annual operating time (h)
$C_L$ = Production worker hourly rate (USD $h^{-1}$)
$EQ_r$= Equity ratio
$C_D$= Total debt costs (USD)
$D_r$= debt ratio (%)
$C_{TEQ}$= Total equity costs (USD)
$f_{CRD}$= Capital recovery factor for debt
$f_{CREQ}$= Capital recovery factor for equity
$D_p$= Annual debt payment (USD)
$EQ_p$= Annual equity recovery (USD)
$C_{cap}$= Minimum annual cost of capital expenditures (USD)
$C_{total}$ = Total minimum annual costs (USD)

<u>Equation list</u>

All cost values are in United States dollar amounts (USD).

Equation 1. Time of batch

$$t_b = t_{gf} + t_m$$

Equation 2. Total number of cells in a single bioreactor after maturation

$$N_c = F_c B_V$$

Equation 3. Total volume occupied by cells

$$V = N_c V_c$$

Equation 4. Cell mass in bioreactor per batch

$$M_b = V \rho_c$$

Equation 5. Annual ACBM production per bioreactor

$$M_{BY} = M_b \, b_{BY}$$

Equation 6. Bioreactors needed to match desired annual beef production

$$B_T = \frac{M_{DY}}{M_{BY}}$$

Equation 7. Equipment costs equation

$$C_{eq} = \sum_j f_{Aj} \, C_{Uj} \left(\frac{U_{aj}}{U_j}\right)^{f_s}$$

Equation 8. Fixed equipment costs

$$C_F = f_L \, C_{eq}$$

Equation 9. Fixed manufacturing costs

$$C_{FM} = f_{FM} \, C_F$$

Equation 10. Minimum annual operating costs

$$C_{op} = C_{FM} + C_{mY} + C_{O_2Y} + C_E \, E_{Hm} + C_E \, E_{BR} + C_E \, E_{ACBMR} + C_{Lab} + C_W$$

Equation 11. Cells in bioreactor during growth phase

$$c_t = 2^{\frac{t}{t_D}} c_o$$

Equation 12. Glucose consumption rate during growth phase

$$\frac{dGCR_B}{dt} = GCR_c \times c_t$$

Equation 13. Total glucose required for growth phase per ACBM batch

$$G_{Gg} = \int_{t=0}^{t=t_{gf}} GCR_B \, dt$$

Equation 14. Total glucose required for maturation phase per ACBM batch

$$G_{GM} = GCR_B \times t_m$$

Equation 15. Total glucose required per batch

$$M_G = G_{Gg} + G_{GM}$$

Equation 16. Total required media charges per batch

$$m_{ch} = G_G/G_{Gch}$$

Equation 17. Total media volume required per batch

$$V_b = m_{ch} V_{ch}$$

Equation 18. Total media volume per year

$$V_m = V_b b_y$$

Equation 19. Total annual costs of media

$$C_{mY} = V_m C_{mL}$$

Equation 20. Oxygen uptake rate

$$OUR_B = OTR_B = kA(e_{con} - a_{con})$$

Equation 21. Initial oxygen in the for the system

$$O_2^i = \frac{V_b \times \rho_m \times P_{O_2}}{O_2^{mol}}$$

Equation 22. Oxygen uptake rate changing with time

$$\frac{dOUR_B}{dt} = OUR_c \times c$$

Equation 23. Total oxygen required for growth phase per ACBM batch

$$O_2^g = \int_{t=0}^{t=t_{gf}} OUR_B \, dt$$

Equation 24. Total oxygen required for maturation phase per ACBM batch

$$O_2^M = OUR_B \times t_m$$

Equation 25. Total oxygen required per ACBM batch

$$O_2^b = O_2^i + O_2^g + O_2^M$$

Equation 26. Total amount of oxygen required per year

$$O_2 = O_2^b b_y$$

Equation 27. Total annual costs of oxygen

$$C_{O_{2Y}} = O_2 C_{O_2}$$

Equation 28. Estimation of energy to heat media to required temperature

$$E_{Hm} = \frac{M_{mY} \times \Delta T \times W_{C_v}}{\in_{Hm}}$$

Equation 29. Glucose combustion reaction

$$C_6H_{12}O_6 + 6\ O_2 \rightarrow 6CO_2 + 6\ H_2O + \text{heat}$$

Equation 30. Estimation of energy usage for bioreactor cooling per ACBM batch

$$E_{BR} = \frac{O_2 \times h}{\in_{BR}}$$

Equation 31. Estimation of annual energy usage for cooling of ACBM

$$E_{ACBMR} = \frac{M_{DY} \times \Delta T \times ACBM_{C_v}}{\in_{ACBMR}}$$

Equation 32. Cost of energy per kWh from public supplier

$$C_{EP} = 0.0969 C_{NG} + 6.78$$

Equation 33. Cost of self-generated electric/energy per kWh from a boiler-turbine system

$$C_{bT} = \frac{C_{NGP}}{\epsilon_{bT}}$$

Equation 34. Cost of energy per kWh

$$C_E = f_{EP} C_{EP} + f_{bT} C_{bT}$$

Equation 35. Annual process water and wastewater costs

$$C_W = V_m C_{PW} + V_m C_{WF} + V_m C_{BO}$$

Equation 36. Required manpower for operation

$$P = \sum_{j}^{N} P_j$$

Equation 37. Labor cost correction factor

$$f_{lab} = f_C f_{Sca} f_T f_Q f_B f_O$$

Equation 38. Estimated annual labor costs

$$C_{Lab} = t_y f_{lab} C_L P$$

Equation 39. Equity ratio

$$EQ_r = 100\% - D_r$$

Equation 40. Total debt costs

$$C_D = C_F D_r$$

Equation 41. Total equity costs

$$C_{TEQ} = EQ_r \, C_F$$

Equation 42. Capital recovery factor for debt

$$f_{CRD} = I_D(1 + I_D)^{L_e} / \ ((1 + I_D)^{L_e - 1})$$

Equation 43. Capital recovery factor for equity

$$f_{CREQ} = I_{EQ}(1 + I_{EQ})^{L_e} / ((1 + I_{EQ})^{L_e - 1})$$

Equation 44. Annual debt payment

$$D_p = f_{CRD} C_D$$

Equation 45. Annual equity recovery

$$EQ_p = f_{CREq}C_{TEq}$$

Equation 46. Minimum annual cost of capital expenditures

$$C_{cap} = D_p + Eq_p$$

Equation 47. Total minimum annual cost

$$C_{total} = C_{cap} + C_{op}$$

**Fig. S1. Costs comparison of the average United States industrial electricity and natural gas (USD $kWh^{-1}$)1999-2019**

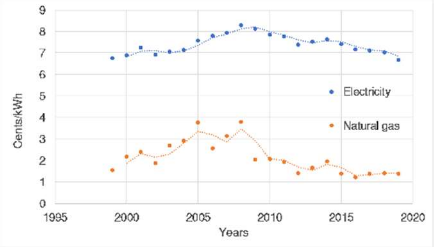


Costs comparison of the average United States industrial electricity and natural gas (USD $kWh^{-1}$) 1999-2019. Information was obtained from the United States EIA and average costs were normalized to January 2019 US currency(*33*, *34*).

**Fig. S2. Linear relationship between electricity and natural gas cost.**

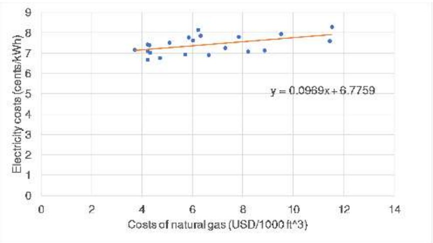


Linear relationship between electricity and natural gas cost. This relationship was used to determine equation 32. Information was obtained from the United States EIA and average costs were normalized to January 2019 US currency(*33*, *34*).

### Table S1a. Model variable inputs: Operations

| Scenarios | inoculum concentration (cells/ml) | Inoculum bioreactor volume (L) | Seed bioreactor volume (L) | Seed bioreactor (cell/ml) | Bioreactor volume ($m^3$) | Desired and achievable cell concentration (cell/ml) | Desired mass of meat produced (kg) |
|---|---|---|---|---|---|---|---|
| 1 | $1.00x10^{7}$ | 2.00 | $2.00x10^{2}$ | $1.00x10^{7}$ | $2.00x10^{1}$ | $1.00x10^{7}$ | $1.21x10^{8}$ |
| 2 | $9.50x10^{7}$ | 2.00 | $2.00x10^{2}$ | $9.50x10^{7}$ | $2.00x10^{1}$ | $9.50x10^{7}$ | $1.21x10^{8}$ |
| 3 | $9.50x10^{7}$ | 2.00 | $2.00x10^{2}$ | $9.50x10^{7}$ | $2.00x10^{1}$ | $9.50x10^{7}$ | $1.21x10^{8}$ |
| 4 | $2.00x10^{8}$ | 2.00 | $2.00x10^{2}$ | $2.00x10^{8}$ | $2.00x10^{1}$ | $2.00x10^{8}$ | $1.21x10^{8}$ |

### Table S1a. Model variable inputs: Operations

| Scenarios | Adjusted value factor for bioreactor | Lang factor | Maturation time (h) | Annual operating time (h) | Bioreactor scale factor | Fixed manufacturing costs factor | Bioreactor unit costs (USD/$m^3$) |
|---|---|---|---|---|---|---|---|
| 1 | 1.29 | 2.00 | 240.00 | 8,760.00 | 0.60 | 0.15 | $5.00x10^{4}$ |
| 2 | 1.29 | 2.00 | 156.00 | 8,760.00 | 0.60 | 0.15 | $5.00x10^{4}$ |
| 3 | 1.29 | 2.00 | 156.00 | 8,760.00 | 0.60 | 0.15 | $5.00x10^{4}$ |
| 4 | 1.29 | 2.00 | 24.00 | 8,760.00 | 0.60 | 0.15 | $5.00x10^{4}$ |

### Table S1b. Model variable inputs: Cell attributes

| Scenarios | Average single cell volume ($m^3$/ cell) | Average single cell density (kg/$m^3$) | Hours per doubling (h) | Glucose consumption rate per cell (mol/h cell) | Rate of oxygen consumption per cell (mol/h cell) |
|---|---|---|---|---|---|
| 1 | $5.00x10^{-15}$ | $1.06x10^{3}$ | 24 | $4.13x10^{-13}$ | 1.80E-14 |
| 2 | $5.00x10^{-15}$ | $1.06x10^{3}$ | 16 | $2.07x10^{-13}$ | 1.80E-14 |
| 3 | $5.00x10^{-15}$ | $1.06x10^{3}$ | 16 | $2.07x10^{-13}$ | 1.80E-14 |
| 4 | $5.00x10^{-15}$ | $1.06x10^{3}$ | 8 | $4.13x10^{-14}$ | 1.80E-14 |

### Table S1c. Model variable inputs: Media

| Scenarios | Basal media (USD/l) | Ascorbic acid 2-phosphate (g/L) | Ascorbic acid 2-phosphate (USD/g) | NAHCO3 (g/L) | NAHCO3 (USD/g) | Sodium selenite (g/L) | Sodium selenite (USD/g) |
|---|---|---|---|---|---|---|---|
| 1 | 3.12 | $6.40x10^{-2}$ | 7.84 | $5.43x10^{-1}$ | 0.01 | $1.40x10^{-5}$ | 0.10 |
| 2 | 3.12 | $6.40x10^{-2}$ | 7.84 | $5.43x10^{-1}$ | 0.01 | $1.40x10^{-5}$ | 0.10 |
| 3 | 3.12 | $6.40x10^{-2}$ | 7.84 | $5.43x10^{-1}$ | 0.01 | $1.40x10^{-5}$ | 0.10 |
| 4 | 0.24 | $6.40x10^{-2}$ | 0.00 | $5.43x10^{-1}$ | 0.00 | $1.40x10^{-5}$ | 0.00 |

**Table S1c. Model variable inputs: Media continued 1**

| Scenarios | Insulin (g/L) | Insulin (USD/g) | Transferrin (g/L) | Transferrin (USD/g) | FGF-2 (g/L) | FGF-2 (USD/g) | TGF-b§ (g/L) | TGF-b§ (USD/g) |
|---|---|---|---|---|---|---|---|---|
| 1 | $1.94x10^{2}$ | 340.00 | $1.07x10^{2}$ | 400.00 | $1.00x10^{-4}$ | $2.01x10^{6}$ | $2.00x10^{-6}$ | $8.09x10^{7}$ |
| 2 | $1.94x10^{2}$ | 340.00 | $1.07x10^{2}$ | 400.00 | $5.00x10^{-5}$ | $1.00x10^{6}$ | $2.00x10^{-6}$ | $8.09x10^{7}$ |
| 3 | $1.94x10^{2}$ | 340.00 | $1.07x10^{2}$ | 400.00 | $5.00x10^{-5}$ | 0.00 | $2.00x10^{-6}$ | $8.09x10^{7}$ |
| 4 | $1.94x10^{2}$ | 0.00 | $1.07x10^{2}$ | 0.00 | 0.00 | 0.00 | $2.00x10^{-6}$ | $0.00 |

**Table S1c. Model variable inputs: Media continued 2**

| Scenarios | Percentage of oxygen in initial charge (w/w) | Oxygen (USD/ton) | Glucose (mol/l) | Density of media (kg/l) |
|---|---|---|---|---|
| 1 | 2.00 | $4.00x10^{1}$ | $1.78x10^{-2}$ | 1.00 |
| 2 | 2.00 | $4.00x10^{1}$ | $2.67x10^{-2}$ | 1.00 |
| 3 | 2.00 | $4.00x10^{1}$ | $2.67x10^{-2}$ | 1.00 |
| 4 | 2.00 | $4.00x10^{1}$ | $3.56x10^{-2}$ | 1.00 |

**Table S1d. Model variable inputs: Utility**

| Scenarios | Boiler energy efficiency (%) | Percentage of electricity self-generated (%) | Temperature of water/media entering facility (˚C) | Desired Temperature of media entering bioreactor (˚C) | Specific heat of water (kWh/ kg (˚C)) | Energy efficiency of media heating system (%) | Heat released per mol of oxygen consumed (kWh) | Energy efficiency of bioreactor cooling system (%) |
|---|---|---|---|---|---|---|---|---|
| 1 | 85 | 50 | 20 | 37 | $1.16x10^{-3}$ | 100 | $1.30x10^{-1}$ | 100 |
| 2 | 85 | 50 | 20 | 37 | $1.16x10^{-3}$ | 100 | $1.30x10^{-1}$ | 100 |
| 3 | 85 | 50 | 20 | 37 | $1.16x10^{-3}$ | 100 | $1.30x10^{-1}$ | 100 |
| 4 | 85 | 50 | 20 | 37 | $1.16x10^{-3}$ | 100 | $1.30x10^{-1}$ | 100 |

**Table S1d. Model variable inputs: Utility continued**

| Scenarios | Specific heat of ACBM (kWh/kg ˚C) | Temperature of ACBM in bioreactor (˚C) | Temperature of cooled ACBM (˚C) | Energy efficiency of ACBM cooling system (%) | natural gas cost (dollars per 1000 ft$^{3}$) | Natural gas (cents per kWh) | Process water cost (USD/m$^{3}$) | Wastewater filtration treatment costs (USD/m$^{3}$) | Biological oxidation of wastewater costs (USD/m$^{3}$) |
|---|---|---|---|---|---|---|---|---|---|
| 1 | $6.22x10^{-4}$ | 37 | 4 | 100 | 4.17 | 1.42 | 0.63 | 0.51 | 0.57 |
| 2 | $6.22x10^{-4}$ | 37 | 4 | 100 | 4.17 | $1.42 | 0.63 | 0.51 | 0.57 |
| 3 | $6.22x10^{-4}$ | 37 | 4 | 100 | 4.17 | $1.42 | 0.63 | 0.51 | 0.57 |
| 4 | $6.22x10^{-4}$ | 37 | 4 | 100 | 4.17 | $1.42 | 0.63 | 0.51 | 0.57 |

**Table S1e. Model variable inputs: Labor**

| Scenarios | Production worker hourly rate (USD/h) | Country effect | Supervising and clerical assistance | Advanced technology and automating | Skilled and qualified level of the personnel | Social benefits | Overtime work | Bioreactors labor factor |
|---|---|---|---|---|---|---|---|---|
| 1 | 13.68 | 1.00 | 1.20 | 0.80 | 1.50 | 1.40 | 1.25 | 1.00 |
| 2 | 13.68 | 1.00 | 1.20 | 0.80 | 1.50 | 1.40 | 1.25 | 1.00 |
| 3 | 13.68 | 1.00 | 1.20 | 0.80 | 1.50 | 1.40 | 1.25 | 1.00 |
| 4 | 13.68 | 1.00 | 1.20 | 0.80 | 1.50 | 1.40 | 1.25 | 1.00 |

**Table S1f. Model variable inputs: Finance**

| Scenarios | Debt ratio (%) | Interest rate on Debt (%/y) | Economic life (y) | Interest cost of equity (%/y) |
|---|---|---|---|---|
| 1 | 90 | 5 | 20.00 | 15 |
| 2 | 90 | 5 | 20.00 | 15 |
| 3 | 90 | 5 | 20.00 | 15 |
| 4 | 90 | 5 | 20.00 | 15 |

Model variable inputs. Inputs without unit in parentheses are unitless.

**Table S2. Sensitivity analysis numerical results**

| Algorithm | Average single cell density (rho_c) | Average single cell volume (V_c) | Glucose concentration (conc_glu) | Glucose consumption rate per cell (GCR_c) | FGF-2 cost (C_fgf2) | FGF-2 concentration (conc_fgf2) | Maturation time (t_m) | TGF-b concentration (conc_tgfb) | Oxygen consumption rate per cell (OUR_c) |
|---|---|---|---|---|---|---|---|---|---|
| DGSM | $6.83x10^{3}$ | $1.00x10^{0}$ | $2.70x10^{-2}$ | $5.70x10^{-1}$ | $2.40\ x10^{-3}$ | $5.07x10^{-2}$ | $8.03x10^{-3}$ | $4.93x10^{-2}$ | $8.68x10^{-2}$ |
| SSA | $1.00x10^{0}$ | $9.66x10^{-1}$ | $9.48x10^{-1}$ | $8.80x10^{-1}$ | $8.50x10^{-1}$ | $7.47x10^{-1}$ | $6.95x10^{-1}$ | $2.16x10^{-3}$ | $1.69x10^{-3}$ |
| DMIM | $8.90x10^{-1}$ | $1.00x10^{0}$ | $9.47x10^{-1}$ | $7.58x10^{-1}$ | $7.83x10^{-1}$ | $9.10x10^{-1}$ | $5.98x10^{-1}$ | $1.37x10^{-2}$ | $5.13x10^{-2}$ |
| FAST | $7.82x10^{-1}$ | $1.00x10^{0}$ | $5.83x10^{-1}$ | $8.63x10^{-1}$ | $4.97x10^{-1}$ | $8.50x10^{-1}$ | $6.94x10^{-1}$ | $1.59x10^{-4}$ | $1.93x10^{-6}$ |
| MM | $1.00x10^{0}$ | $9.70x10^{-1}$ | $9.91x10^{-1}$ | $9.53x10^{-1}$ | $9.11x10^{-1}$ | $9.09x10^{-1}$ | $8.62x10^{-1}$ | $1.44x10^{-2}$ | $1.44x10^{-8}$ |
| RBD-FAST | $1.00x10^{0}$ | $7.94x10^{-1}$ | $9.96x10^{-1}$ | $7.54x10^{-1}$ | $7.86x10^{-1}$ | $7.11x10^{-1}$ | $8.22x10^{-1}$ | $1.39x10^{-1}$ | $7.48x10^{-2}$ |

Sensitivity analysis numerical results. DGSM = Derivative-based Global Sensitivity Measure, SSA = Sobol Sensitivity Analysis, DMIM = Delta Moment-Independent Measure, FAST = Fourier Amplitude Sensitivity Analysis MM = Morris Method and RBD-FAST = Random Balance Designs-Fourier Amplitude Sensitivity Test. This analysis was performed using peer reviewed open source SALib Python package for this work (*20*).

**Table S3. Potential industrial scale equipment for ACBM production.**

| Equipment | Unit | Unit costs ($1000's) | Scale index | Production Operators required (P) | Adjusted value factor ($f_{Aj}$) | Accounted for in equipment cost analysis |
|---|---|---|---|---|---|---|
| Centrifugal pumps | Power (kW) | 5 | 0.60 | 0.1 | 1.42 | - |
| Plate filters | Area ($m^2$) | 3 | 0.75 | 1.0 | 1.64 | - |
| Media holding vessel | Volume ($m^3$) | 10 | 0.50 | 0.2 | 1.29 | - |
| Heat exchanger | Area ($m^2$) | 3 | 0.65 | 0.5 | 1.29 | - |
| Inoculum bioreactor | Volume ($m^3$) | 50 | 0.60 | 1.0 | 1.29 | - |
| Seed bioreactor | Volume ($m^3$) | 50 | 0.60 | 1.0 | 1.29 | - |
| Bioreactors | Volume ($m^3$) | 50 | 0.60 | 1.0 | 1.29 | + |
| Positive displacement pump | Power (kW) | 5 | 0.60 | 0.1 | 1.42 | - |

Potential industrial scale equipment for ACBM production. Created using information from *Food Plant Economics* and CEPI (*35–37*).

**Table S4. Annual United States national industrial grid electricity costs 1999-2019**

| Year | Average nominal consumer cost per year (cents $kWh^{-1}$) | Inflation adjusted cost (cents $kWh^{-1}$) |
|---|---|---|
| 1999 | 4.42 | 6.77 |
| 2000 | 4.63 | 6.9 |
| 2001 | 5.04 | 7.25 |
| 2002 | 4.88 | 6.94 |
| 2003 | 5.11 | 7.08 |
| 2004 | 5.25 | 7.14 |
| 2005 | 5.72 | 7.59 |
| 2006 | 6.15 | 7.81 |
| 2007 | 6.39 | 7.95 |
| 2008 | 6.95 | 8.29 |
| 2009 | 6.83 | 8.14 |
| 2010 | 6.76 | 7.85 |
| 2011 | 6.81 | 7.78 |
| 2012 | 6.66 | 7.4 |
| 2013 | 6.88 | 7.52 |
| 2014 | 7.09 | 7.63 |
| 2015 | 6.90 | 7.43 |
| 2016 | 6.75 | 7.17 |
| 2017 | 6.87 | 7.12 |
| 2018 | 6.92 | 7.03 |

Annual United States industrial national grid electricity costs 1999-2019. Information was obtained from the United States EIA and average costs were normalized to January 2019 US currency(*33*, *34*).

**Table S5. Annual United States national industrial natural gas costs 1999-2019**

| Year | Average nominal cost per year (USD thousand cubic feet$^{-1}$) | Inflation adjusted cost (cents kWh$^{-1}$) |
|---|---|---|
| 1999 | 3.08 | 1.55 |
| 2000 | 4.45 | 2.19 |
| 2001 | 5.08 | 2.40 |
| 2002 | 4.02 | 1.88 |
| 2003 | 5.91 | 2.70 |
| 2004 | 6.51 | 2.92 |
| 2005 | 8.67 | 3.77 |
| 2006 | 7.82 | 2.58 |
| 2007 | 7.65 | 3.13 |
| 2008 | 9.66 | 3.79 |
| 2009 | 5.23 | 2.05 |
| 2010 | 5.44 | 2.08 |
| 2011 | 5.12 | 1.93 |
| 2012 | 3.85 | 1.41 |
| 2013 | 4.64 | 1.67 |
| 2014 | 5.58 | 1.98 |
| 2015 | 3.91 | 1.39 |
| 2016 | 3.49 | 1.22 |
| 2017 | 4.08 | 1.39 |
| 2018 | 4.17 | 1.42 |

Annual United States national average natural gas costs 1999-2019. Information was obtained from the United States EIA and average costs were normalized to January 2019 US currency(*33*, *34*).

**Table S6. Cost of process and wastewater treatment**

| Utility | Cost (USD $m^{-3}$) |
|---|---|
| Process water | 0.63 |
| Wastewater filtration treatment | 0.51 |
| Biological oxidation of wastewater | 0.57 |

Cost of process and wastewater treatment. Cost were reported in *Food Plant Economics* and were adjusted to account for inflation reported in January 2019 US currency (*34*, *37*).

16. S. Parulekar, G. Birol, A. Cinar, C. Undey, in *Batch Fermentation Modeling: Monitoring*

*and Control* (CRC Press, ed. 1, 2003), vol. 93, pp. 1–19.

26. M. D. McKay, R. J. Beckman, W. J. Conover, A comparison of three methods for

selecting values of input variables in the analysis of output from a computer code. *Technometrics*. **21**, 239 (1979).
